# Extensive Mammalian Germline Genome Engineering

**DOI:** 10.1101/2019.12.17.876862

**Authors:** Yanan Yue, Yinan Kan, Weihong Xu, Hong-Ye Zhao, Yixuan Zhou, Xiaobin Song, Jiajia Wu, Juan Xiong, Dharmendra Goswami, Meng Yang, Lydia Lamriben, Mengyuan Xu, Qi Zhang, Yu Luo, Jianxiong Guo, Shengyi Mao, Deling Jiao, Tien Dat Nguyen, Zhuo Li, Jacob V. Layer, Malin Li, Violette Paragas, Michele E. Youd, Zhongquan Sun, Yuan Ding, Weilin Wang, Hongwei Dou, Lingling Song, Xueqiong Wang, Lei Le, Xin Fang, Haydy George, Ranjith Anand, Shi Yun Wang, William F. Westlin, Marc Güell, James Markmann, Wenning Qin, Yangbin Gao, Hong-jiang Wei, George M. Church, Luhan Yang

## Abstract

Xenotransplantation, specifically the use of porcine organs for human transplantation, has long been sought after as an alternative for patients suffering from organ failure. However, clinical application of this approach has been impeded by two main hurdles: 1) risk of transmission of porcine endogenous retroviruses (PERVs) and 2) molecular incompatibilities between donor pigs and humans which culminate in rejection of the graft. We previously demonstrated that all 25 copies of the PERV elements in the pig genome could be inactivated and live pigs successfully generated. In this study, we improved the scale of porcine germline editing from targeting a single repetitive locus with CRISPR to engineering 13 different genes using multiple genome engineering methods. we engineered the pig genome at 42 alleles using CRISPR-Cas9 and transposon and produced PERVKO·3KO·9TG pigs which carry PERV inactivation, xeno-antigen KO and 9 effective human transgenes. The engineered pigs exhibit normal physiology, fertility, and germline transmission of the edited alleles. *In vitro* assays demonstrated that these pigs gain significant resistance to human humoral and cell mediated damage, and coagulation dysregulations, similar to that of allotransplantation. Successful creation of PERVKO·3KO·9TG pigs represents a significant step forward towards safe and effective porcine xenotransplantation, which also represents a synthetic biology accomplishment of engineering novel functions in a living organism.

**One Sentence Summary:** Extensive genome engineering is applied to modify pigs for safe and immune compatible organs for human transplantation

## Introduction

The limited availability of organs for human transplantation is a serious unmet medical need. Owing to their similar organ sizes and physiology, pigs have long been thought to be a promising source of organs, tissues, and cells for human transplantation. However, although many attempts have been made over the years, pig-to-human transplantation has been impeded by two major hurdles: 1) risk of cross-species transmission of the porcine endogenous retroviruses (PERVs) and 2) molecular incompatibilities between the porcine graft and the human recipient (*1, 2*).

PERVs can infect human cells *in vitro*, propagate among cells, and recombine to adapt to human host, which may pose a zoonosis concern for pig-to-human transplantation (*2, 3*). Integration of PERV into human genome can potentially lead to immunodeficiency and tumorigenesis (*4*). Because PERV sequences are a part of the porcine genome, they cannot be eliminated by bio-secure breeding as could be done for other zoonotic pathogens. We have previously demonstrated the genome-wide inactivation of all 25 copies of the PERV elements by using CRISPR to target repetitive elements (hereinafter referred to as “PERVKO”) and successful production of PERV-free pigs, eliminating the potential risk of viral transmission (*3*).

Another barrier of clinical xenotransplantation is organ rejection due to molecular incompatibility between pig organs and human immune system. In particular, some glycan epitopes, primarily α-Gal (galactose-alpha-1,3-galactose), Neu5Gc (N-glycolylneuraminic acid), and the SDa epitope, are unique to pigs and binding of preformed antibodies initiates hyperacute graft rejection through the activation of the complement cascade (*5-8*). Genetic inactivation of GGTA1 (alpha-1,3-galactosyltransferase), CMAH (cytidine monophosphate-N-acetylneuraminic acid hydroxylase), and B4GALNT2 (beta-1,4-N-acetyl-galactosaminyltransferase 2) removes α-Gal, Neu5Gc, and the SDa epitopes from cell surface, which has been shown to attenuate this rapid graft rejection (*9-13*).

In addition, overexpression of human complement regulatory proteins, including CD46 (membrane cofactor protein), CD55 (decay-accelerating factor), and CD59 (MAC-inhibitory protein), could further reduce hyperacute rejection and prolong graft survival (*14-17*). Beyond humoral injury and hyperacute reaction, acute cellular reactions to the xenograft, including the infiltration of Natural Killer (NK) cells and macrophages, contribute to xenotransplantation rejection (*18*). It has been demonstrated that expression of human B2M-HLA-E fusion protein in porcine endothelial cells reduces NK-mediated cell toxicity through inhibitory signaling (*19*), and expression of human CD47 reduces macrophage-mediated toxicity through the interaction of CD47 with SIRPα (*20*).

In addition, incompatibilities between the pig and human coagulation systems could also lead to thrombotic microangiopathy and systemic consumptive coagulopathy observed in many xenotransplantation preclinical experiments (*1*). In particular, porcine TFPI (tissue factor pathway inhibitor) does not effectively inhibit human factor Xa and VIIa/TF (tissue factor) complex to prevent thrombin formation (*21*). Also, porcine thrombomodulin (THBD) is incapable of binding to human thrombin to activate antithrombotic protein C to prevent clot formation (*22*). Beyond that, porcine CD39 is inactivated upon endothelial cell activation and its expression is insufficient to inhibit human platelet aggregation (*21, 23*). To address these incompatibilities, several genetic modifications targeting the coagulation pathway have been tested and coagulation compatibility was restored with some success (*24-26*).

To date, more than 40 genetic modifications have been attempted either individually or in combination on pigs with the goal to mitigate incompatibility-related hyperacute rejection, delayed xenograft rejection, and acute cellular rejection (*14*). Pigs carrying triple knockout of GGTA1, CMAH, and B4GALNT2 (hereinafter referred to as “3KO”) have been created, with their peripheral blood mononuclear cells showing minimal xenoreactive antibody binding compared to pigs carrying single knockout of GGTA1 or double knockout of GGTA1 and CMAH (*14, 27*). In addition, various human transgenes have been tested, some of which demonstrated beneficial outcome in preclinical models (*14*). Encouragingly, one recent study showed that a cardiac xenograft from the pig carrying knockout of GGTA1 and overexpression of hCD46 and hTHBD achieved life-supporting function in baboon for 6 months (*28*). Another study showed that a rhesus macaque monkey lived for more than 400 days with a renal xenograft from the pig carrying knockout of GGTA1 and overexpression of hCD55 (*29*). Furthermore, porcine islet cell xenotransplantation in diabetic nonhuman primates (NHPs) showed significantly improved engraftment rate using islets from pigs carrying knockout of GGTA1 (*30*). These studies support that long-term graft survival is achievable using genetically engineered xenograft organs.

Despites years of efforts, no pig organs have been engineered with the combined feature of PERV elimination, improved immunological and coagulation compatibility, which is ideal for clinical applications (*14*).

In this study, we sought to use different genome modification tools and cloning technology to create clinically usable pig organs.

## Results

We first engineered 3KO·9TG pigs carrying 3KO to eliminate major xeno-antigens and 9 transgenes (TG) with the goal to enhance the immunological and coagulation compatibility between pigs and humans. Next, we inactivated all PERVs from the 3KO·9TG pig genome to produce PERVKO·3KO·9TG pigs carrying 3KO, 9TG, and PERVKO (Fig. S1).

To engineer 3KO·9TG pigs, we electroporated wild-type porcine ear fibroblasts with CRISPR-Cas9 reagents targeting the GGTA1, CMAH, and B4GALNT2 genes and plasmids encoding PiggyBac transposase (*31*) and carrying a transgenic construct consisting of the nine human transgenes (hCD46, hCD55, hCD59, hB2M, hHLA-E, hCD47, hTHBD, hTFPI, and hCD39) (Fig. 1A, Methods). Next, we isolated and expanded single-cell clones, and screened for clones carrying the desired 3KO mutations by Sanger sequencing and whole genome sequencing (Fig. 1B). In addition, we validated the presence of 9TG by conventional PCR (Fig. 1C). After obtaining the modified cells, we performed somatic cell nuclear transfer (SCNT) using the validated clones and successfully produced 3KO·9TG pigs.

**Fig. 1.**
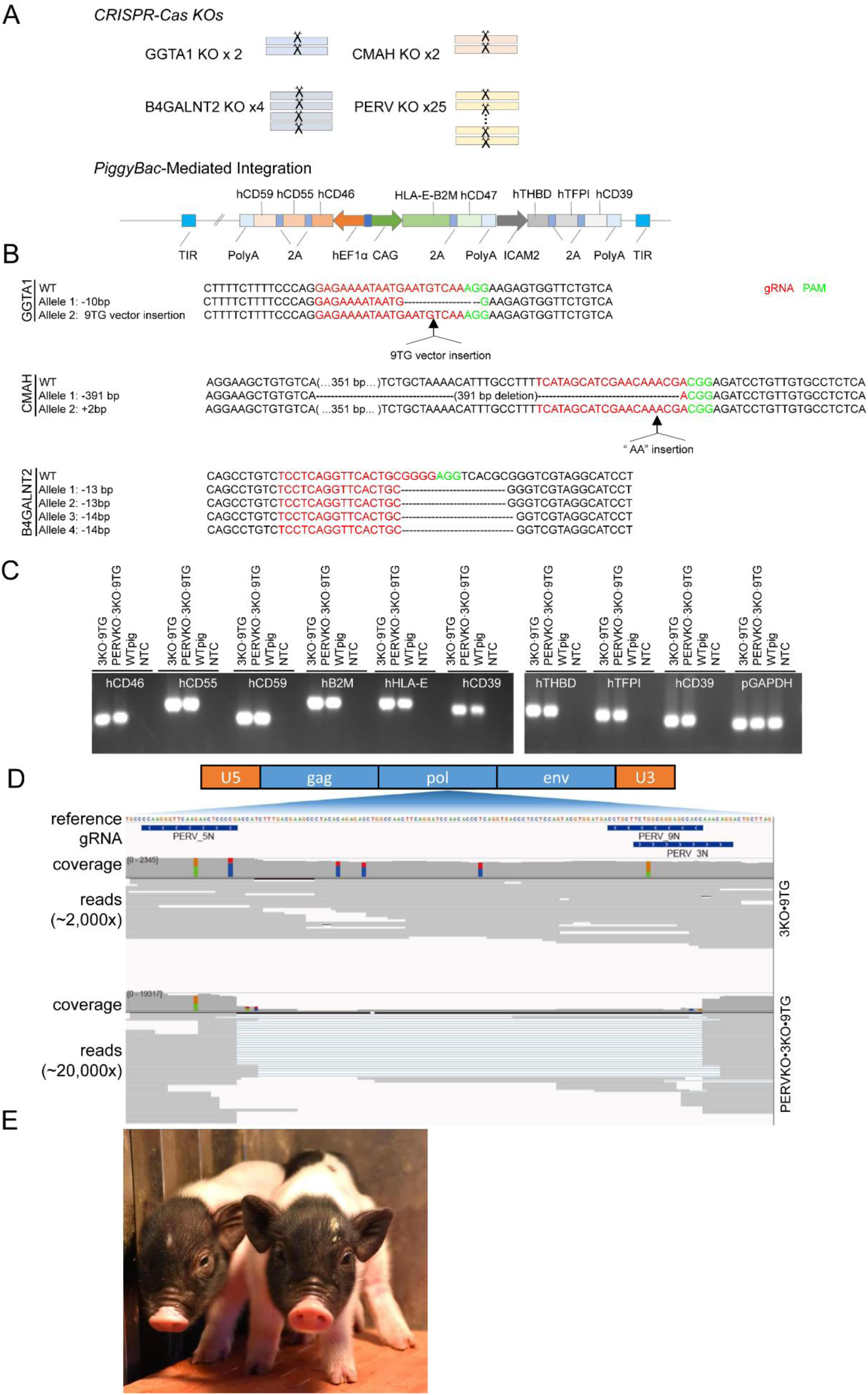
PERVKO·3KO·9TG pig engineering and validation of the 3KO, 9TG and PERVKO edits at the genomic level. (A) Schematic diagram of the 42 modified alleles. We generated 3KO and PERVKO using CRISPR-Cas9 with gRNAs targeting the 2 copies of GGTA1, 2 copies of CMAH, 4 copies of B4GALNT2, and 25 copies of PERVs. We generated 9TG using PiggyBac-mediated random integration of the 9 human transgenes into the pig genome. The transgenes are expressed from 3 transcription cassettes, with each cassette expressing two to three genes linked by the porcine teschovirus 2A (P2A) peptide. TIR: Terminal Inverted Repeats of PiggyBac transposon; hEF1α1/CAG/ICAM2: promoters; PolyA: Polyadenylation signal. **(B)** WGS confirmation of the frameshift mutations of GGTA1, CMAH and B4GALNT2 in 3KO·9TG pigs and PERVKO·3KO·9TG pigs. The GGTA1 locus was found to carry a 10 bp deletion in one allele and an insertion of the transgenic construct in the other allele. CMAH was impacted by a 391 bp deletion in one allele and a 2 bp insertion into the other allele. B4GALNT2 carries deletions of 13 bps, 13 bps, 14 bps, and 14 bps respectively for the four alleles. (C) Agarose gel image of PCR products suggests the presence of the 9 human transgenes in the 3KO·9TG and PERVKO·3KO·9TG fetal fibroblasts, and the absence in the WT fetal fibroblasts and no template controls. Primers used are listed in Table S1. (D) Deep capture sequencing confirmed the complete knockout of PERVs. The raw reads for 3KO·9TG pigs (∼2,000X) and PERVKO·3KO·9TG pigs (∼20,000X) are shown below a schematic PERV gene structure. Reads are grouped by their sequence composition and shown proportionally according to their sequencing depth. The vertical lines in red, blue, green, and orange in the coverage track represent single nucleotide change from reference allele to T, C, A, G respectively. (E) Image of two 5-day-old PERVKO·3KO·9TG piglets.

To generate PERVKO·3KO·9TG pigs, we electroporated the 3KO·9TG pig fibroblasts with CRISPR-Cas9 reagents targeting the reverse transcriptase (*pol*) gene common to all 25 copies of the PERV elements (*3*). Next, we screened and selected single-cell clones carrying exclusively large deletions encompassing the catalytic core of the *pol* gene as measured by deep sequencing (Fig. 1D). Finally, we selected the set of clones based on the presentation of a normal karyotype and with these clones, successfully produced PERVKO·3KO·9TG pigs via SCNT (Fig. 1E and S2).

We next sought to examine closely the on-target and off-target effects of genetic modifications in PERVKO·3KO·9TG pigs. To this end, we performed 10X whole genome sequencing on WT, 3KO·9TG, and PERVKO·3KO·9TG ear fibroblasts. Consistent with our deep sequencing data, we confirmed that mutations introduced to the 25 copies of the PERV elements and 8 alleles of the 3KO genes are frameshift insertions or deletions (Fig. 1D and 1B). In addition, we confirmed the presence of all nine transgenes in the porcine genome (Fig. 1C). We did not observe any difference in structural variants between WT and 3KO·9TG pigs, or between 3KO·9TG pigs and PERVKO·3KO·9TG pigs, indicating gross genomic stability for these engineered pigs. For the small indels, we examined all 1,211 predicted off-target sites and found two small insertions in the B4GALNT2 gRNA off-target sites in 3KO·9TG pigs compared to WT; however, neither affect protein coding sequence (Fig. S3). Comparing PERVKO·3KO·9TG genome to 3KO·9TG genome, we also found two deletions and one insertion within two PERV gRNA off-target sites (Fig. S3). Neither of these is located within any annotated protein coding region. Of note, we could not rule out the possibility that these mutations are derived from spontaneous somatic mutations (*32, 33*). Given the lack of functional implications and together with normal *in vivo* pathophysiology data of engineered pigs (Fig. S4), we conclude that the germline engineered PERVKO·3KO·9TG pigs maintained genomic stability.

Having confirmed the genomic modifications at DNA level, we further examined if PERVKO·3KO·9TG pigs have the proper 3KO and 9TG expression. We first performed RNA-seq on fibroblasts and endothelial cells, and found that both 3KO·9TG pigs and PERVKO·3KO·9TG pigs expressed all transgenes at levels comparable to or higher than that from human umbilical vein endothelial cells (HUVECs) (Fig. 2A). In addition, we observed comparable transgene expression profile and level in both pig umbilical vein endothelial cells (PUVECs) and fibroblasts, suggesting that the transgenes are ubiquitously expressed among these cell types. We next characterized protein expression in the engineered pigs. We observed diminished glycan markers of α-Gal, Neu5Gc, and SDa on cell surface, which suggests functional elimination of the 3 genes responsible for synthesizing these glycan epitopes in both 3KO·9TG pigs and PERVKO·3KO·9TG pigs (Fig. 2B). By FACS analysis of PUVECs, we observed that both 3KO·9TG pigs and PERVKO·3KO·9TG pigs express all transgenes at the protein level. Indeed, eight out of the nine transgenes are robustly expressed at a level comparable to that of HUVECs. Intriguingly, THBD expression is detectable but at a lower level. Consistent with FACS analysis, we detected that PERVKO·3KO·9TG pigs lack the three glycan antigens (Fig. 2C) and express the eight transgenes in the kidneys, with THBD close to background level (Fig. 2C). Taken together, we conclude that our 3KO and 9 TG genetic modifications largely translate into successful RNA and protein expression at the cellular and tissue level in engineered pigs.

**Fig. 2.**
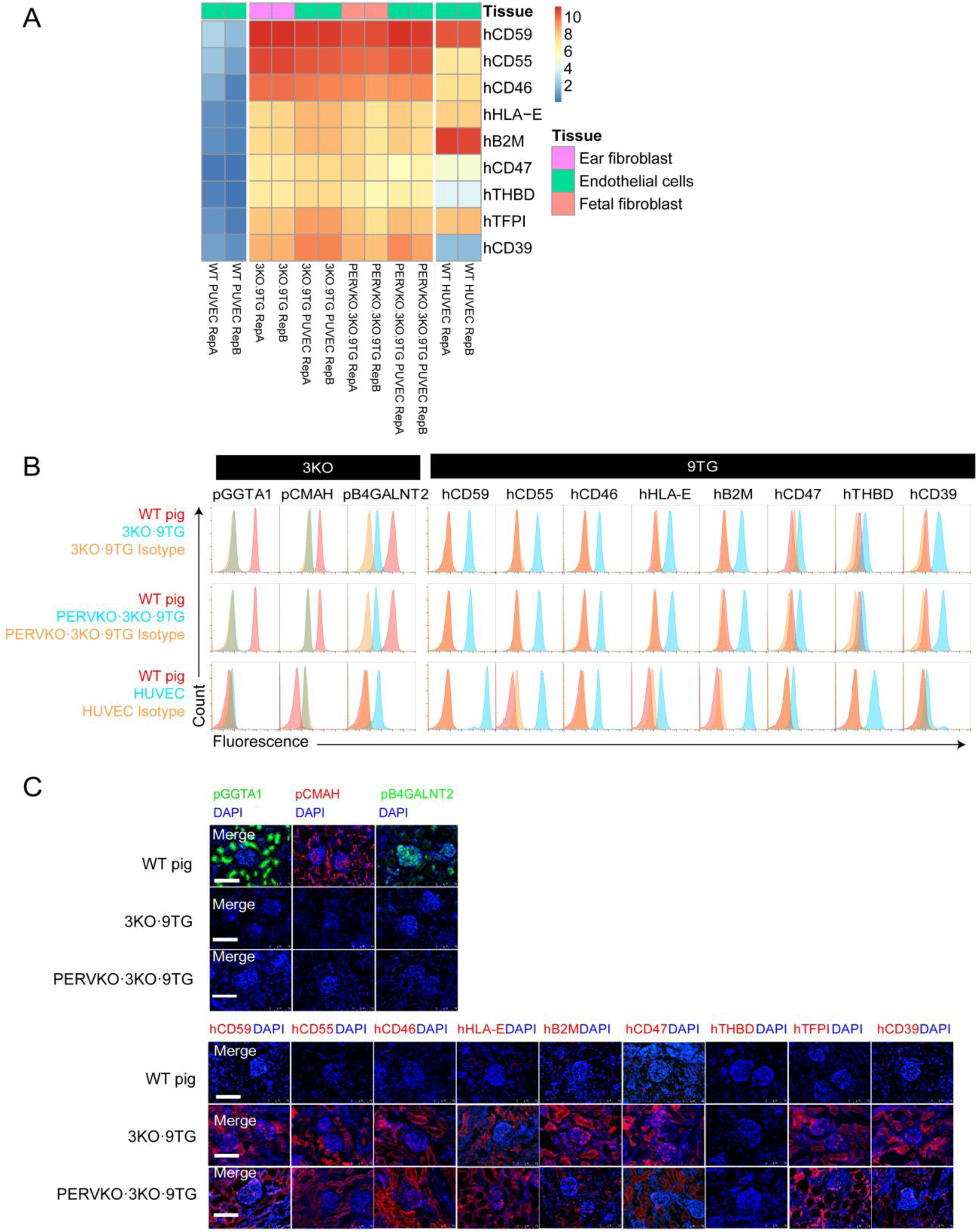
Validation of 3KO and 9TG in 3KO·9TG pigs and PERVKO·3KO·9TG pigs at mRNA and protein level. (A) Heatmap of transgene expression measured by RNA-Seq in WT PUVECs, 3KO·9TG PUVECs and ear fibroblasts, PERVKO·3KO·9TG PUVECs and fetal fibroblasts, and WT HUVECs. Each row represents one transgene and each column represents one sample. Expression level is color coded in blue-yellow-red to represent low-medium-high. The color bar on top of the heatmap indicate the sample type. (B) FACS validation of 3KO and 9TG. 3KO·9TG PUVECs and PERVKO·3KO·9TG PUVECs show comparable TG protein expression as HUVECs, with the exceptions of hCD39 (higher) and hTHBD (lower). (C) Immunofluorescence staining validation of 3KO and 9TG in the 3KO·9TG and PERVKO·3KO·9TG kidney cryosections. Antibodies used are listed in Table S2.

To assess the overall fitness of the engineered pigs, we examined the physiology, fertility, and transmission of the genetic modifications of the engineered pigs to the offspring. We observed that both PERVKO pigs and 3KO·9TG pigs, although having been extensively engineered on the PERV loci, immunological and coagulation pathways, show normal blood cell counts, including total white blood cell and platelet, monocyte, neutrophil, and eosinophil counts (Fig. S4A). We also observed normal vital organ functions for liver, heart, and kidney of engineered pigs (Fig. S4B-4D). In addition, engineered pigs have similar prothrombin and thrombin time as compared with WT pigs (Fig. 4E), suggesting normal coagulation functions.

In addition, we found PERVKO pigs and 3KO·9TG pigs are fertile and produce a normal average litter size of seven. The offspring from breeding PERVKO pigs with WT pigs carry ∼50% PERV inactivated alleles in their liver, kidney, and heart tissues, indicating that the knockout alleles are stably inherited following Mendelian genetics (Fig. S5). Similarly, all the offspring of 3KO·9TG pigs and WT pigs are heterozygous for 3KO and approximately half of the them carry 9TG (Fig. S6A), with expression validated at both the mRNA (Fig. S6B) and protein level (Fig. S6C). This suggests that the genetic modifications are stable and are not silenced at the F1 generation. Therefore, we conclude that the engineered PERVKO pigs and 3KO·9TG pigs exhibit normal physiology, fertility, and germline transmission of the edited alleles.

With the engineered pigs, we examined if the genetically modified pigs acquired novel functions as designed. We first tested if the genetic modifications allow the modified pig cells to evade preformed human antibody binding. Both 3KO·9TG PUVECs and PERVKO·3KO·9TG PUVECs exhibited approximately 90% reduction in antibody binding from human IgG and IgM when compared to WT PUVECs, confirming that the antibody barrier to xenotransplantation can be greatly mitigated by 3KO (Fig. 3A-3C). In addition, when incubated with human complement from pooled human sera, PERVKO·3KO·9TG PUVECs with 3KO and co-expressing human complement modulators hCD46, hCD55, and hCD59 demonstrated minimal *in vitro* human complement toxicity, similar to their HUVEC counterpart (Fig. 3D).

**Fig. 3.**
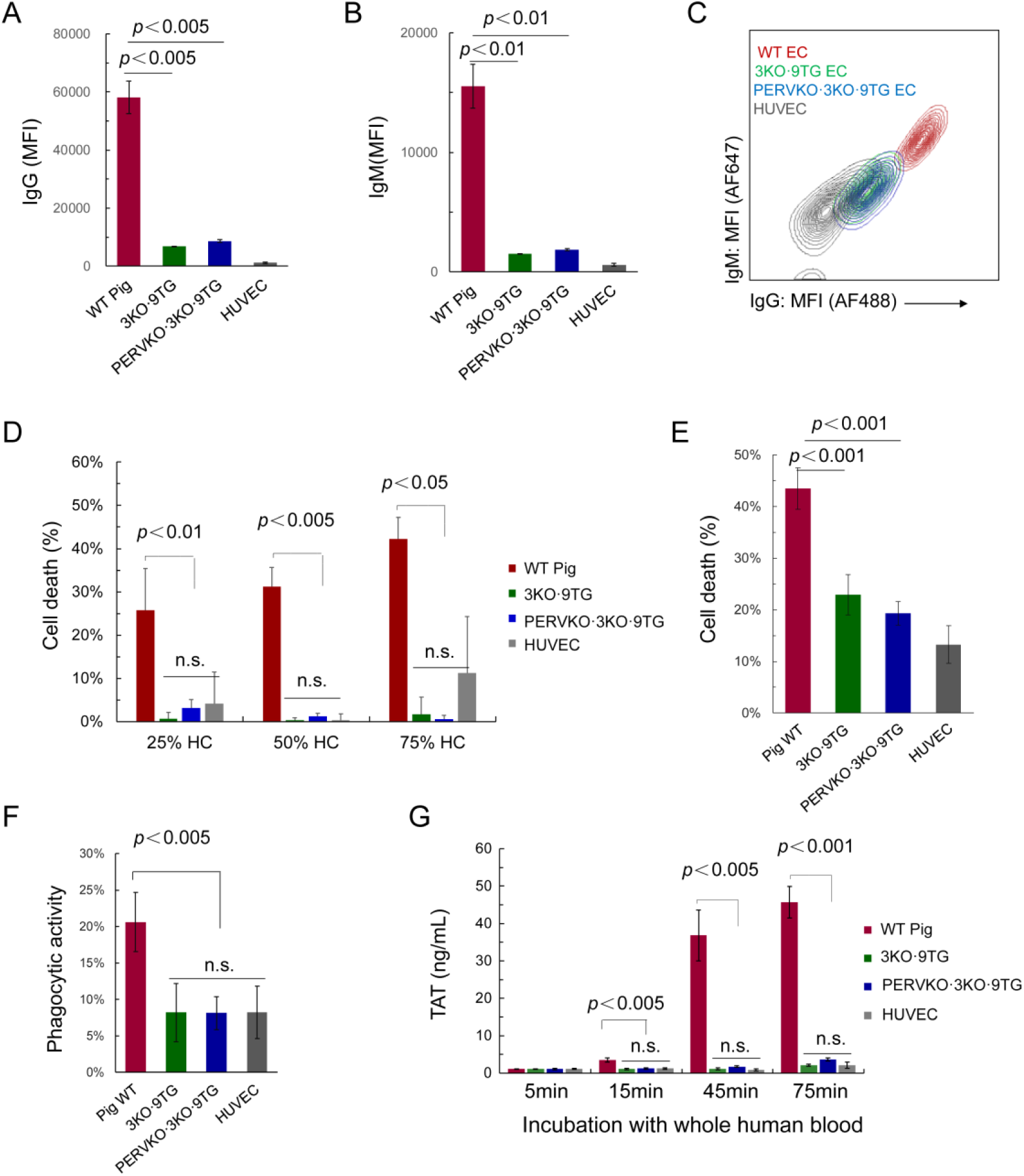
Functional validation of PERVKO·3KO·9TG pigs in mitigating human antibody binding, complement toxicity, NK cell toxicity, and modulating coagulation function. (A-C) 3KO·9TG PUVECs and PERVKO·3KO·9TG PUVECs show substantially less binding of human IgG and IgM as compared to their WT counterpart. Antibody binding of pooled human serum to PUVECs and HUVECs (positive control) was measured by FACS (n = 3). Hereinafter error bars indicate standard deviation. (D) 3KO·9TG PUVECs and PERVKO·3KO·9TG PUVECs show comparable antibody-dependent complement cytotoxicity as HUVECs and significantly lower than WT PUVECs (n = 4). Hereinafter “n.s.” denotes no statistical significance (*P* > 0.05) among the 3KO·9TG PUVEC, PERVKO·3KO·9TG PUVEC and HUVEC groups by pairwise T-test analysis. (E) 3KO·9TG PUVECs and PERVKO·3KO·9TG PUVECs show significantly lower NK-mediated cytotoxicity compared to their WT counterpart (n = 4). (F) 3KO·9TG PUVECs and PERVKO·3KO·9TG PUVECs reveal reduced phagocytosis by a human macrophage cell line (n = 4). (G) 3KO·9TG PUVECs and PERVKO·3KO·9TG PUVECs mediate very low level of thrombin-antithrombin (TAT) formation upon incubation with whole human blood for the indicated time (n = 4), comparable to HUVECs and significantly lower than WT PUVECs.

Second, we examined if PERVKO·3KO·9TG pigs are more resistant to injury mediated by human innate cellular immunity. Consistent with previous reports (*19, 34*), PERVKO·3KO·9TG PUVECs expressing hHLA-E/hB2M demonstrated significantly higher resistance to NK-mediated cell killing compared with that of WT PUVECs (Fig. 3E). In addition, PERVKO·3KO·9TG pigs expressing hCD47 exhibited elevated suppression of human macrophage phagocytosis, potentially via the CD47-SIRPα signaling pathway (Fig. 3F) (*20*). Taken together, these results suggest PERVKO·3KO·9TG pig xenograft has been successfully engineered and is expected to be more resistant to attack by human innate cell immunity.

Lastly, we examined if PERVKO·3KO·9TG pigs can attenuate the dysregulated activation of platelets and coagulation cascades. When vascularized WT porcine organs are transplanted, preformed human antibodies, complement, and innate immune cells can induce endothelial cell activation and trigger coagulation and inflammation (*1*). The incompatibility between coagulation regulatory factors from pig endothelial cells and human blood leads to abnormal platelet activation and thrombin formation (*1, 21*), exacerbating the damage. To address this issue, we overexpressed hCD39 in PERVKO·3KO·9TG pigs. The ADPase function of CD39 hydrolyzes ADP, a potent platelet antagonist, into AMP, which inhibits human platelet activation and aggregation. *In vitro* ADPase biochemical assay showed significantly higher CD39 activity in PERVKO·3KO·9TG PUVECs when compared with WT PUVECs and HUVECs, consistent with its higher mRNA and protein expression (Fig. S7). In addition, molecular incompatibilities of coagulation regulators (e.g., TFPI) between pig and human render the extrinsic coagulation regulation ineffective. To address this issue, we overexpressed hTFPI in PERVKO·3KO·9TG pigs, which translocate to cell surface following endothelial cell activation (*35*). As expected, activated PERVKO·3KO·9TG PUVECs effectively bind and neutralize human Xa, which should mitigate coagulation and reduce the formation of thrombin-antithrombin (TAT) complex (Fig. S8). To examine coagulation reaction holistically, we co-cultured human whole blood with PERVKO·3KO·9TG PUVECs and found minimal TAT formation, similar to that of HUVECs (Fig. 3G), suggesting that PERVKO·3KO·9TG pigs acquired enhanced coagulation compatibility with human factors.

Collectively, our results indicate that PERVKO·3KO·9TG pigs acquired enhanced compatibility with the human immune system with attenuated human antibody binding, complement toxicity, NK cell toxicity, phagocytosis, and restored coagulation regulation.

## Discussion

Genetically engineered pigs hold great promise in addressing the unmet medical need of organs for human transplantation. In this report, we use different engineering tools and created PERVKO·3KO·9TG pigs with 13 genes and 42 alleles modified to eradicate PERV activity and enhance human immunological and coagulation compatibility. Extensive analysis showed that the engineered pig cells exhibit reduced human antibody binding, complement toxicity, NK cell toxicity, and coagulation dysregulation. We also examined and validated the normal pathophysiology, fertility, and genetic inheritability of our engineered pigs. The successful production of PERVKO·3KO·9TG pigs has brought us one step closer in using pigs to produce safe and effective organs for clinical transplantation.

Our work demonstrated the feasibility of repurposing the pig immunological markup for human transplantation with extensive genome editing. Work is in progress to test the safety and effectiveness of the Pig 3.0 organs in non-human primate to understand to what degree we could dampen the immunological barrier through engineered organs.

We also identified some future improvement opportunities. For example, compared with other genes, THBD protein is expressed at a lower level. We are investigating several possibilities, including transgene isoform choice, post-transcriptional and post-translational modifications, and cellular localization in the porcine host.

More importantly, successful generation of PERVKO·3KO·9TG pigs demonstrates the power of synthetic biology to extensively engineer the mammalian germline to confer novel functions in mammals. In PERVKO·3KO·9TG pigs, we use the combination of CRISPR and transposon to deleted 25 copies of the PERV elements, 8 alleles of the xenogeneic genes, and concurrently expressed 8 human transgenes to physiologically relevant levels. It extends the record of genome modifications to 13 different genes in large animal models. We also identified some future improvement opportunities. For example, compared with other genes, THBD protein is expressed at a lower level. We are investigating into several possibilities, including transgene isoform choice, post-transcriptional and post-translational modifications, and cellular localization in the porcine host. With the ability to execute complex genetic engineering at this scale, we are in a position to engineer additional modifications and functions in pigs for therapeutic purpose. We envision PERVKO·3KO·9TG pigs can be further genetically engineered to achieve additional novel functions, such as immune tolerance, organ longevity, and viral immunity.

## Acknowledgments

The pig cloning work was supported by National Key R&D Program of China (grant number: 2019YFA0110700). We are indebted to Dr. Geoffrey Yang of Harvard University for the careful review of our manuscript. We wish to thank Dr. Qizhi Tang and Dr. Phil O’Conner for their excellent advices, Dr. Yang Yang and Dr. Qing Yang from Third Affiliated Hospital of Sun Yat-sen University, Dr. Hongbo Liu from Henan Chuangyuan Biotechnology Co. Ltd., and colleagues at Qihan Bio Inc. and eGenesis Inc. for their technical assistance and helpful discussions.

## Author contributions

Y.L., G.M.C., Y.G. and W.Q. envisioned and supervised the whole project; W.J. and H.Z. supervised pig cloning and production; Y.Y., Y.K. and W.X. designed the experiments and wrote the manuscript; Y.Y., Y.K., Y.Z., X.S., L. Lamriben, J.W., J.X., M.X., Q.Z., Y.L., J.V.L., M.L., V.P., M.E. Y, Z.S., Y.D., W.W., H.D., L.S., X.W., L.Le, X.F, H.G., R.A. and S.Y.W. performed experiments; W.X., D.G., M.Y. and M.G. analyzed the data; J.G., S.M., D. J., T.D.N. and Z.L. performed pig cloning and generated pigs; and J.M. and W.F.W. revised the manuscript.

