## Supplementary materials for "Extensive Mammalian Germline Genome Engineering"

**This PDF file includes:**

Materials and Methods
Figs. S1 to S8
Tables S1 to S2

### Materials and Methods

CRISPR-Cas9 gRNA design

We used the R library DECIPHER to design specific gRNAs (PERV-3N: 5'-TCTGGCGGGAGCCACCAAAC-3', PERV-5N: 5'-GGCTTCGTCAAAGATGGTCG-3', PERV-9N: 5'-TTCTAAGCAGTCCTGTTTGG-3') to target specifically all *pol* catalytic sequences in the 3KO·9TG genome as described before (*1*). In addition, we used specific gRNAs to target GGTA1, CMAH and B4GALNT2, respectively (GGTA1: 5'-GCTGCTTGTCTCAACTGTAA-3', CMAH: 5'-GAAGCTGCCAATCTCAAGGA-3', B4GALTN2: 5'- GATGCCCGAAGGCGTCACAT-3').

Cell culture

Porcine fetal fibroblast cells and ear fibroblast cells were maintained in Dulbecco's modified Eagle's medium (DMEM, Invitrogen) high glucose with sodium pyruvate supplemented with 20% fetal bovine serum (Invitrogen), 100 U/mL penicillin/streptomycin (Pen/Strep, Invitrogen) and 10 mM HEPES (Thermo Fisher Scientific). All cells were maintained in a humidified tri-gas incubator at 37°C and 5% $CO_2$, 90% $N_2$, and 5% $O_2$. Porcine umbilical vein endothelial cells (PUVECs) were freshly isolated and cultured in Prigrow II medium (abm) supplemented with 10% fetal bovine serum (Gibco), 100 U/mL penicillin/streptomycin (Pen/Strep, Invitrogen) and 10mM HEPES (Thermo Fisher Scientific). Human umbilical vein endothelial cells (HUVECs, ATCC, PCS-100-010) were cultured in vascular cell basal medium supplemented with Endothelial Cell Growth Kit-BBE (ECG kit, ATCC). Human NK-92 cell line was cultured in Minimum Essential Medium Alpha (α-MEM, Gibco) supplemented with 12.5% fetal bovine serum (Gibco), 12.5% fetal equine serum (FES, Solarbio) and 100 U/mL penicillin/streptomycin (Pen/Strep, Invitrogen). The human macrophage cell line THP-1 was cultured in RPMI 1640 (BI) supplemented with 10% fetal bovine serum (Gibco) and 100 U/mL penicillin/streptomycin (Pen/Strep, Invitrogen). Differentiation of THP-1 cells was performed in 62.5 nM Phorbol-12-myristate-13-acetate (PMA, Sigma) for 3 days and confirmed by attachment of these cells to tissue-culture plastic.

3KO·9TG cell production

To generate 3KO·9TG ("3KO" denotes the triple knockout of porcine GGTA1, CMAH, and B4GALNT2 genes, while "9TG" denotes the expression of nine human transgenes, including hCD46, hCD55, hCD59, hTHBD, hTFPI, hCD39, hB2M, hHLA-E and hCD47) cells, we transfected 1 million porcine fetal fibroblasts with 2.5 µg Cas9 expression plasmid, 1.5 µg B4GALNT2, 1.0 µg CMAH and 0.5 µg GGTA gRNA expression plasmids, 1 µg Super PiggyBac plasmid (Cat. # PB210PA-1, System Biosciences) and 4 µg transgene expression plasmid. Nine days after transfection, cells were stained with Isolectin B4 (GGTA1, ALX-650-001F-MC05, Enzo), CD46 (A15776, Invitrogen) and CD39 (560239, BD Biosciences) antibodies. Subsequently, cells that are negative for GGTA1 and positive for both CD46 and CD39 were sorted by FACS into 96-well plates. When individual clones reached 30-50% confluency, cells were transferred and expanded whereas aliquots were used to genotype using KAPA mouse genotyping kit (KR0385, KAPA Biosystems). The 3KO was genotyped using NGS and the presence of the 9TG was confirmed by PCR using primers listed in Table S1.

**Supplementary Table 1. Summary of PCR primers used in 3KO and 9TG genotyping**

| Primer Name | Sequence (5' to 3') |
|---|---|
| GGTA1_NGSF | **ACACTCTTTCCCTACACGACGCTCTTCCGATCT**CGCTCGTTGACTATTCATC |
| GGTA1_NGSR | **GACTGGAGTTCAGACGTGTGCTCTTCCGATCT**ACTAGGAGATTAGAGGAGACT |
| B4GALNT2_NGSF | **ACACTCTTTCCCTACACGACGCTCTTCCGATCT**TCCCAATCTGTGATCTTTGA |
| B4GALNT2_NGSR | **GACTGGAGTTCAGACGTGTGCTCTTCCGATCT**CACCCTCGGGAATGAGTA |
| CMAH_NGSF | **ACACTCTTTCCCTACACGACGCTCTTCCGATCT**GAGGGAGGGCTTTCAAAC |
| CMAH_NGSR | **GACTGGAGTTCAGACGTGTGCTCTTCCGATCT**TCAGGCGGCTCTTATTCT |
| hCD46_F | ACCTGGTAGAAGGGAGTGTC |
| hCD46_R | TTGGTGGTTCTTCACAAGCAT |
| hCD55_F | CATCTGTTCCAGCAGCACTC |
| hCD55_R | AGAGCAGGTTGAGCATTAGGA |
| hCD59_F | GGTATCCAAGGTGGTTCTGTTC |
| hCD59_R | ACAGTCAGCAGTAGGGTTGG |
| hB2M_F | CACTCTGACCTGTCTTTCTCTA |
| hB2M_R | CATGTCTCTGTCCCATTTAACA |
| hHLAE_F | TGGTATCATTGCTGGTCTTGTC |
| hHLAE_R | CACTCCCTTGAGCTGAATCTG |
| hCD47_F | ATCCTGGCTCTGGCTCAATT |
| hCD47_R | AGCATTCAGAGGTTCTTCAACA |
| hTHBD_F | GTTAGGTGTTCTGGTTCTTGGT |
| hTHBD_R | GAGAAGCATTCAGGAAGGTAGC |
| hTFPI_F | GGCTCCTGCTCCTCTGAAT |
| hTFPI_R | TTACAAGGACCATCATCAGCTT |
| hCD39_F | CTGGCTATCCTGGGTTTCTCT |
| hCD39_R | AACCAGCATCCAGAACAATACC |
| pGAPDH_F | CCGCGATCTAATGTTCTCTTTC |
| pGAPDH_R | TTCACTCCGACCTTCACCAT |

PERVKO·3KO·9TG cell production

The 3KO·9TG fibroblasts were edited for generating PERVKO·3KO·9TG as previously described (*1*). We also conducted genotyping for the clones derived from the sorted single cells. The procedure for genotyping was as described previously (*2*). Briefly, we sorted single cells into 96-well PCR plates with each well carrying a 5 µL lysis mixture, which contained 0.5 µL 10×KAPA express extract buffer (KAPA Biosystems), 0.1 µL of 1U/µL KAPA Express Extract Enzyme and 4.4 µL water. We incubated the lysis reaction at 75ºC for 15 min and inactivated the reaction at 95ºC for 5 min. All reactions were then added to 20 µL PCR reactions containing 1× KAPA 2G fast (KAPA Biosystems) and 0.2 µM PERV Illumina primers as follows,

Illumina_PERV_*pol* forward:
5'-ACACTCTTTCCCTACACGACGCTCTTCCGATCTCGACTGCCCCAAGGGTTCAA-3'
Illumina_PERV_*pol* reverse:
5'-GTGACTGGAGTTCAGACGTGTGCTCTTCCGATCTTCTCTCCTGCAAATCTGGGCC-3'.

Reactions were incubated at 95ºC for 3 min followed by 30 (for single cell) or 25 (for single cell clones) cycles of 95ºC, 20 s; 59ºC, 20 s and 72ºC, 10 s. To add the Illumina sequence adaptors, 3µL of reaction products were then added to 20 µL of PCR mix containing 1×KAPA 2G fast (KAPA Biosystems) and 0.3 µM primers carrying Illumina sequence adaptors. Reactions were incubated at 95ºC for 3 min, followed by 20 (for single cell) or 10 (for single cell clones) cycles of 95ºC, 20 s; 59ºC, 20 s and 72ºC, 10 s. PCR products were examined on EX 2% gels (G401002, Invitrogen), followed by the recovery of ~360 bp target products from the gel. These products were then mixed at roughly the same amount, purified (28115, Qiagen), and sequenced with MiSeq Personal Sequencer (Illumina). We then analyzed deep sequencing data and determined the PERV editing efficiency using CRISPR-CAS.

<u>Somatic cell microinjection to produce SCNT embryos and embryo transfer for pig cloning</u>

The somatic cell microinjection procedure was reported previously (*3*). All animal experiments were approved by the Animal Care Committee of Yunnan Agricultural University, China. All chemicals were purchased from Sigma Chemical Co. (St. Louis, MO, USA), unless otherwise indicated. Porcine ovaries were collected from Hongteng Abattoir (Chenggong Ruide Food Co., Ltd, Kunming, Yunnan Province, China). The ovaries were transported to the laboratory at 25°C to 30°C in 0.9% (w/v) NaCl solution supplemented with 75 mg/mL potassium penicillin G and 50 mg/mL streptomycin sulfate. The cumulus cell-oocyte complexes (COCs) were isolated from the follicles of 3-6 mm in diameter, and then cultured in 200 µL TCM-199 medium supplemented with 0.1 mg/mL pyruvic acid, 0.1 mg/mL L-cysteine hydrochloride monohydrate, 10 ng/mL epidermal growth factor, 10% (v/v) porcine follicular fluid, 75 mg/mL potassium penicillin G, 50 mg/mL streptomycin sulfate, and 10 IU/mL eCG and hCG (Teikoku Zouki Co., Tokyo, Japan) at 38.5°C in a humidified atmosphere with 5% CO2 (APC- 30D, ASTEC, Japan). After 38 to 42 hours in-vitro maturation, the expanded cumulus cells of the COCs were removed by repeat pipetting of the COCs in 0.1% (w/v) hyaluronidase.

SCNT was conducted as previously described (*4*, *5*). Briefly, oocytes extruding the first polar body with intact membrane were cultured in NCSU23 medium supplemented with 0.1 mg/mL demecolcine, 0.05 M sucrose, and 4 mg/mL bovine serum albumin (BSA) for 0.5 to 1 hour for nucleus protrusion. The protruded nucleus was then removed along with the polar body by using a bevelled pipette (approximately 20 µm in diameter) in Tyrode's lactate medium supplemented with 10 µM hydroxyethyl piperazineethanesulfonic acid (HEPES), 0.3% (w/v) polyvinylpyrrolidone, and 10% FBS in the presence of 0.1 mg/mL demecolcine and 5 mg/mL cytochalasin B. Fibroblasts with confirmed genotypes were used as nuclear donors. A single donor cell was injected into the perivitelline space of the enucleated oocyte.

Donor cells were fused with the recipient cytoplasts with a single direct current pulse of 200 V/mm for 20 µs by using an embryonic cell fusion system (ET3, Fujihira Industry Co. Ltd., Tokyo, Japan) in a fusion medium which contains 0.25 M D-sorbic alcohol, 0.05 mM $Mg(C_2H_3O_2)_2$, 20 mg/mL BSA and 0.5 mM HEPES (free acid). The reconstructed embryos were cultured in PZM-3 solution (*5*) for 2 hours to allow nucleus reprogramming and then activated with a single pulse of 150 V/mm for 100 µs in an activation medium containing 0.25 M D-sorbic alcohol, 0.01 mM $Ca(C_2H_3O_2)_2$, 0.05 mM $Mg(C_2H_3O_2)_2$ and 0.1 mg/mL BSA. The activated embryos were then cultured in PZM-3 supplemented with 5 mg/mL cytochalasin B for 2 hours at 38.5°C in humidified atmosphere with 5% $CO_2$, 5% $O_2$, and 90% $N_2$ (APM- 30D for further activation, ASTEC, Japan). Reconstructed embryos were then transferred to new PZM-3 medium and cultured in humidified air with 5% $CO_2$, 5% $O_2$, and 90% $N_2$ at 38.5°C for 2 and 7 days to detect the embryo cleavage and blastocyst development ratios, respectively.

Crossbred (Large White/Landrace Duroc) sows with one birth history were used as the surrogate mothers of the constructed embryos. They were examined for estrus at 9:00 am and 6:00 pm daily. The SCNT embryos cultured for 6 hours after activation were surgically transferred to the oviducts of the surrogates. Pregnancy was examined 23 days after embryo transfer using an ultrasound scanner (HS-101 V, Honda Electonics Co. Ltd., Yamazuka, Japan).

Validation of genetic modification at protein level by FACS

Umbilical vein endothelial cells derived from 3KO·9TG, PERVKO·3KO·9TG and WT pigs were analyzed by FACS to characterize the genetic modifications (3KO and 9TG) at protein level. Cells were harvested, fixed and then stained using corresponding primary and secondary antibodies (Supplementary Table 2), according to manufacturer's instructions. Isotype controls were applied at the same final dilution as the specific primary antibodies. After antibody staining, cells were washed twice, and analyzed by FACS using a CytoFLEX S flow cytometer. 5,000 events were analyzed for each sample by using a Flow Jo software.

Characterization of protein expression by immunofluorescence

Neonatal (3-6 days old) porcine kidney cryosections of WT, 3KO·9TG and PERVKO·3KO·9TG pigs were subject to immunofluorescence to characterize the genetic modification (3KO and 9TG) at tissue level. Cryosections were fixed with ice-cold acetone, blocked and then stained using either one-step direct or two-step indirect immunofluorescence techniques. The primary and secondary antibodies used were summarized in Supplementary Table 2. Nuclear staining was performed using ProLong Gold DAPI (Thermo Fisher, P36931). Sections were imaged using a Leica Fluorescence Microscope and analyzed using ImageJ software. All pictures were taken under the same condition to allow accurate comparison of fluorescence intensities among WT, 3KO·9TG and PERVKO·3KO·9TG cryosections.

### Supplementary Table 2. Summary of Antibodies

| Name | Application* | Catalog No.; Supplier | Dilution fold |
|---|---|---|---|
| FITC Isolectin B4 (GGTA1) | FACS, IF | ALX-650-001F-MC05; Enzo | 100 |
| Chicken anti-Neu5Gc Antibody Kit (CMAH) | FACS, IF | 146901; Biolegend | 200 |
| Fluorescein DOLICHOS BIFLORUS AGGLUTININ (B4GALNT2) | FACS, IF | FL-1031; Vector | 100 |
| FITC Mouse anti-human CD46 (MEM-258) | FACS | 315304; Biolegend | 50 |
| PE Mouse anti-human CD55 | FACS | 555694; BD | 100 |
| APC Mouse anti-human CD59 | FACS | MA1-19463; Invitrogen | 50 |
| APC Mouse anti-human CD39 | FACS | 560239; BD | 50 |
| APC Mouse anti-human THBD | FACS | 564123; BD | 50 |
| FITC Mouse anti-human CD47 | FACS | 556045; BD | 25 |
| PE Mouse anti-human HLA-E | FACS | LS-C106985; LSBio | 30 |
| PE Mouse anti-human B2M | FACS | 316306; Biolegend | 100 |
| Mouse anti-human TFPI | FACS | MA5-26774; Invitrogen | 50 |
| FITC mouse IgG1, k Isotype ctrl | FACS | 555748; BD | 50 |
| PE mouse IgG1, k Isotype ctrl | FACS | 555749; BD | 50 |
| APC mouse IgG1, k Isotype ctrl | FACS | 555751; BD | 50 |
| Rabbit anti-human CD59 | IF | HPA026494; Sigma | 100 |
| Mouse anti-human CD55-PE | IF | 555694; BD | 25 |
| Mouse anti-human CD46 | IF | HM2103; Hycult | 100 |
| Rabbit anti-human HLA-E | IF | 14-9953-82; Invitrogen | 100 |
| Mouse anti-human B2M | IF | MA1-19141; Thermo Fisher | 100 |
| Rabbit anti-human CD47 | IF | Ab226837; Abcam | 100 |
| Rabbit anti-human THBD | IF | HPA002982; Sigma | 100 |
| Rabbit anti-human TFPI | IF | HPA005575; Sigma | 100 |
| Rabbit anti-human CD39 | IF | Ab223842; Abcam | 100 |
| EnVision+System-HRP Labelled Polymer Anti-Rabbit | IF | K400311-2; Dako | Ready-to-use |
| EnVision+System-HRP Labelled Polymer Anti-Mouse | IF | K400111-2; Dako | Ready-to-use |
| Goat Anti-Rabbit IgG Alexa 555 | IF | A-21428; Thermo Fisher | 1000 |
| Goat anti-Chicken Alexa Fluor 568 | IF | A-11041; Thermo Fisher | 500 |
| APC mouse anti-human CD56 | NK assay | 555518; BD | 50 |
| FITC mouse anti-human CD31 | NK assay | 555445; BD | 50 |
| FITC mouse anti-pig CD31 | NK assay | MCA1746F; Bio-rad | 50 |
| CD11b monoclonal antibody (M1/70), APC | Phagocytosis assay | 17-0112-81; Thermo Fisher | 100 |
| Goat anti-Human IgM, Alexa | Antibody | A-21249; Thermo | 200 |

| | | | |
|---|---|---|---|
| Fluor 647 | binding | Fisher | |
| Goat Anti-Human IgG, Alexa Fluor 488 | Antibody binding | A-11013; Thermo Fisher | 200 |

*IF: Immunofluorescence

Human antibody binding to porcine endothelial cells

Binding of human IgG and IgM to the porcine and human endothelial cells were assessed by flow cytometry as described (*6*). Briefly, PUVECs and HUVECs were collected, washed twice and resuspended in staining buffer (PBS containing 1% BSA). Heat-inactivated pooled normal human male AB serum (Innovative Research) were diluted 1:4 in staining buffer. 3KO·9TG PUVECs, PERVKO·3KO·9TG PUVECs, WT PUVECs and HUVECs ($1 \times 10^5$ cells per test) were incubated with diluted human serum for 30 min at room temperature, respectively. Cells were then washed with cold staining buffer and incubated with goat anti-human IgG Alexa Fluor 488 (Invitrogen, A11013, 1:200 dilution) and goat anti-human IgM Alexa Fluor 647 (Invitrogen, A21249, 1:200 dilution) for 30 min at room temperature. After washing with cold staining buffer, cells were resuspended in staining buffer containing 7-AAD (559925, BD Biosciences; 1: 100 dilution) to exclude nonviable cells. Fluorescence was acquired on CytoFLEX S flow cytometer and data were analyzed using FlowJo analysis software. For each sample, 5,000 events were collected in the live cell gate and plotted as specific median fluorescence intensity (MFI) which is generated by "test MFI (IgG or IgM) – control (secondary antibody only) MFI".

Human complement-dependent cytotoxicity assay

3KO·9TG PUVECs, PERVKO·3KO·9TG PUVECs, WT PUVECs and HUVECs were harvested, washed twice with PBS, and resuspended in serum-free culture medium. Cells ($1\times10^5$ cells per test) were incubated with a uniform pool of human serum complement (A113, Quidel) at different concentrations (0%, 25%, 50% and 75%) for 45 min at 37$^o$C and 5% $CO_2$. Afterwards, cells were stained with propidium iodide (P3566, Invitrogen; 1:500 dilution) for 5 min and analyzed using a CytoFLEX S flow cytometer. 5,000 events were collected for each sample, and the percentage of PI positive cells was used as the percentage of cell death.

NK cytotoxicity assay

PUVECs and HUVECs were used as target cells and labeled with anti-pig CD31-FITC antibody (Bio-Rad, MCA1746F) and anti-human CD31-FITC antibody (BD, 555445), respectively. Meanwhile, human NK 92 cells were used as effector cells and labeled with anti-human CD56-APC antibody (BD, 555518). The effector (E) and target cells (T) were cocultured for 4 hours at 37°C and 5% $CO_2$, at an E/T ratio of 3.  Cells were stained with propidium iodide for 5 min and then subjected to FACS analysis. The percentage of PI positive cells in $CD31^+$ gate was used to calculate the percentage of killed target cells.

Phagocytosis Assay

Differentiation of human macrophage cell line THP-1 was achieved by 62.5 nM of phorbol myristate acetate (PMA) for 3 days and confirmed by attachment of these cells to tissue-culture plastic. 3KO·9TG PUVECs, PERVKO·3KO·9TG PUVECs, WT PUVECs and HUVECs (target cells) were stained with the fluorescent dyes 5/6-CFSE (Molecular Probes) according to the manufacturer's protocol. CFSE-labeled target cells were incubated with human differentiated THP-1 cells (effector cells) at E/T ratios of 2:1 for 4 hours at 37ºC. Macrophages were counterstained with anti-human CD11b antibody (Thermo Fisher, 17-0112-81) and phagocytosis of CFSE-labeled targets were measured by FACS. Phagocytic activity was calculated as previously described (*7*).

CD39 biochemical ADPase assay

3KO·9TG PUVECs, PERVKO·3KO·9TG PUVECs, WT PUVECs and HUVECs were seeded at $2\times10^4$ per well in a 96-well plate, 1 day before the assay. Cells were incubated with 500 μM ADP (Chrono-Log Corp, #384) for 30 min at 37°C and 5% $CO_2$. Malachite green (MAK307, Sigma) was added to stop the reaction, and absorbance was measured at 630 nm to determine levels of phosphate generation against the standard curve of $KH_2PO_4$.

TFPI activity and human factor Xa binding assay

To prepare for the assay, cells were treated with 1 μM PMA for 6 hours to promote hTFPI translocation to cell surface of 3KO·9TG and PERVKO·3KO·9TG PUVECs. TFPI activity and human factor Xa binding assay was subsequently performed as previously described (*6*)**.** All assays were performed in quadruplicate.

TAT (Thrombin-antithrombin complex) formation assay

3KO·9TG PUVECs, PERVKO·3KO·9TG PUVECs, WT PUVECs and HUVECs were seeded at $3\times10^5$ per well in 6-well plates. After 1 day, cells were incubated with 1 mL of fresh whole human blood (containing 0.5 U/mL heparin) at 37ºC with gentle shaking. At different indicated time points, blood was aspirated, from which plasma was isolated. TAT content in plasma was measured using a Thrombin-Antithrombin Complex Human ELISA Kit (ab108907, Abcam).

Variant calling from whole genome sequencing data

Paired reads are mapped to the Sus Scrofa 11.1 genome (ftp://ftp.ensembl.org/pub/release-91/fasta/sus_scrofa/dna/) by BWA (v0.7.17-r1188) (*8*). Variants (SNPs and INDELs) are called using GATK (v4.0.7.0) (*9*) following the GATK best practice recommendation (*10*) with the standard filter plus requiring a minimum depth of 10.

*In silico* prediction of on/off-target sites

Genome-wide on-target and off-target sites are predicted using CRISPRSeek (v1.22.1) (*11*) in R (v3.5.0) allowing up to 4-mismatches. The input genome is Sus Scrofa 11.1 (ftp://ftp.ensembl.org/pub/release-91/fasta/sus_scrofa/dna/).

Off-target calling from whole genome sequencing data

Filtered variants from GATK within 10 bp distance to the PAM sites of predicted off-targets by CRISPRSeek (v1.22.1) are called as potential off-target modifications. Variants with allele frequency deviate from the parental line significantly more or less than 0.5 are filtered out using Two-proportion Z-test. The assumption for this test is the chance for both alleles to be simultaneously modified is highly unlikely because off-target mutation introduced by CRISPR-Cas9 genome editing is very rare (*12*).

Functional impact analysis of variants

Regardless a variant is an off-target or germline mutation, it is annotated for sequence change at transcript level and amino acid change at protein level to assess its potential functional impact using VEP (variant effect predictor, v93.3) (*13*). High impact variants are specially selected if they can result in frameshift, start gain/lost, stop gain/lost, splice donor/acceptor shift or splice region changes. Whenever available, the variant will be annotated to indicate whether it impacts principle or alternative transcripts using the APPRIS database (*14*).

Transcription analysis from RNA-Seq

RNA-Seq reads are aligned to the Sus Scrofa 11.1 genome using STAR (v2.6.1a) (*15*) under the splicing-aware mode. The expression level is quantified as TPM (transcripts per million) using Salmon (v0.11.3) (*16*) with both the porcine transcriptome and the nine transgenes as reference transcripts.

PERV knock-out efficiency analysis by deep sequencing

Paired reads are first aligned to the PERV capture target sequence using STAR (v2.6.1a) under the splicing-aware mode, followed by alignment position dependent deduplication by Picard (v2.18.14). Deduped paired reads are then merged into fragments by an in-house script. Merged fragments are then re-aligned to the PERV capture target sequence using STAR (v2.6.1a) under the splicing-aware mode. The output BAM file is then analyzed by an in-house R script (v3.5.0) to digest the alignment pattern to assess the distribution of INDELs within the capture target sequence and derive the knock-out efficiency.

Statistical Analysis

All statistical analyses are performed by R (v3.5.0) and Excel (v2016). A p-value $< 0.05$ is significant unless otherwise specified. When multiple tests are involved simultaneously, a p-value correction is performed following the Benjamini–Hochberg procedure to control the overall false discovery rate (FDR). An FDR$<0.05$ is typically used unless otherwise specified.

**Supplementary Figures**

| | |
|---|---|
| Figure S1 | Schematic diagram of generation process of 3KO·9TG pigs and PERVKO·3KO·9TG pigs |
| Figure S2 | Normal karyotypes of 3KO·9TG pigs and PERVKO·3KO·9TG pigs |
| Figure S3 | Summary of off-target analysis on 3KO·9TG pigs and PERVKO·3KO·9TG pigs |
| Figure S4 | PERVKO pigs and 3KO·9TG pigs show similar pathophysiology, compared with WT pigs in parameters measured, including complete blood count (A), liver function (B), heart function (C), kidney function (D), and coagulation function (E). |
| Figure S5 | The genetic modification of PERVKO can be inherited following Mendelian genetics during natural mating production |
| Figure S6 | The genetic modifications (3KO and 9TG) of 3KO·9TG pigs can be transmitted to the next generation following Mendelian genetics through natural mating production, as validated at genomic DNA (A), mRNA (B) and protein Level (C) |
| Figure S7 | 3KO·9TG PUVECs and PERVKO·3KO·9TG PUVECs show significantly higher CD39 ADPase biochemical activity compared to WT PUVECs and HUVECs |
| Figure S8 | Activated 3KO·9TG PUVECs and PERVKO·3KO·9TG PUVECs express human TFPI on cell surface and show significantly higher binding ability to human Xa compared to WT PUVECs and HUVECs *in vitro* |

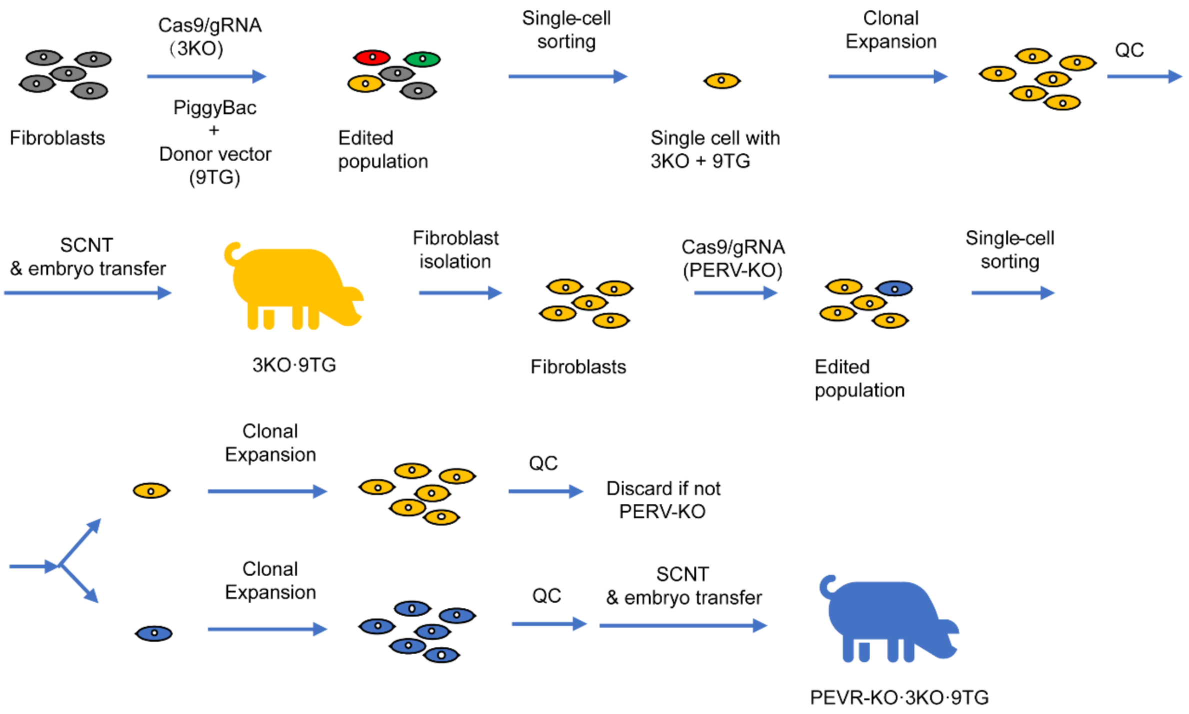


**Fig. S1. Schematic diagram of generation process of 3KO·9TG pigs and PERVKO·3KO·9TG pigs.** Wild type Bama ear fibroblasts were edited using a mixture of Cas9 protein, guide RNAs targeting GGTA1, CMAH and B4GALNT2 genes, as well as a vector carrying the expression cassettes for 9TG (hCD46, hCD55, hCD59, hTHBD, hTFPI, hCD39, hB2M, hHLA-E and hCD47) and a vector encoding PiggyBac transposase. The edited population was sorted into single cell clones carrying 3KO and 9TG integration. The selected clonal cells were used as donors in Somatic Cell Nuclear Transfer (SCNT), reconstructed embryos transferred, and piglets born (designated as 3KO·9TG pigs). In the next round of engineering, fibroblasts from 3KO·9TG pigs were isolated and further edited to inactivate PERVs using Cas9 and guide RNAs targeting the *pol* gene of PERVs. The edited 3KO·9TG cells were single cell sorted for clones carrying PERVKO in addition to 3KO and 9TG. Cells with all PERV alleles inactivated were used as donors in SCNT, embryos transferred, and piglets born (designated as PERVKO·3KO·9TG pigs).

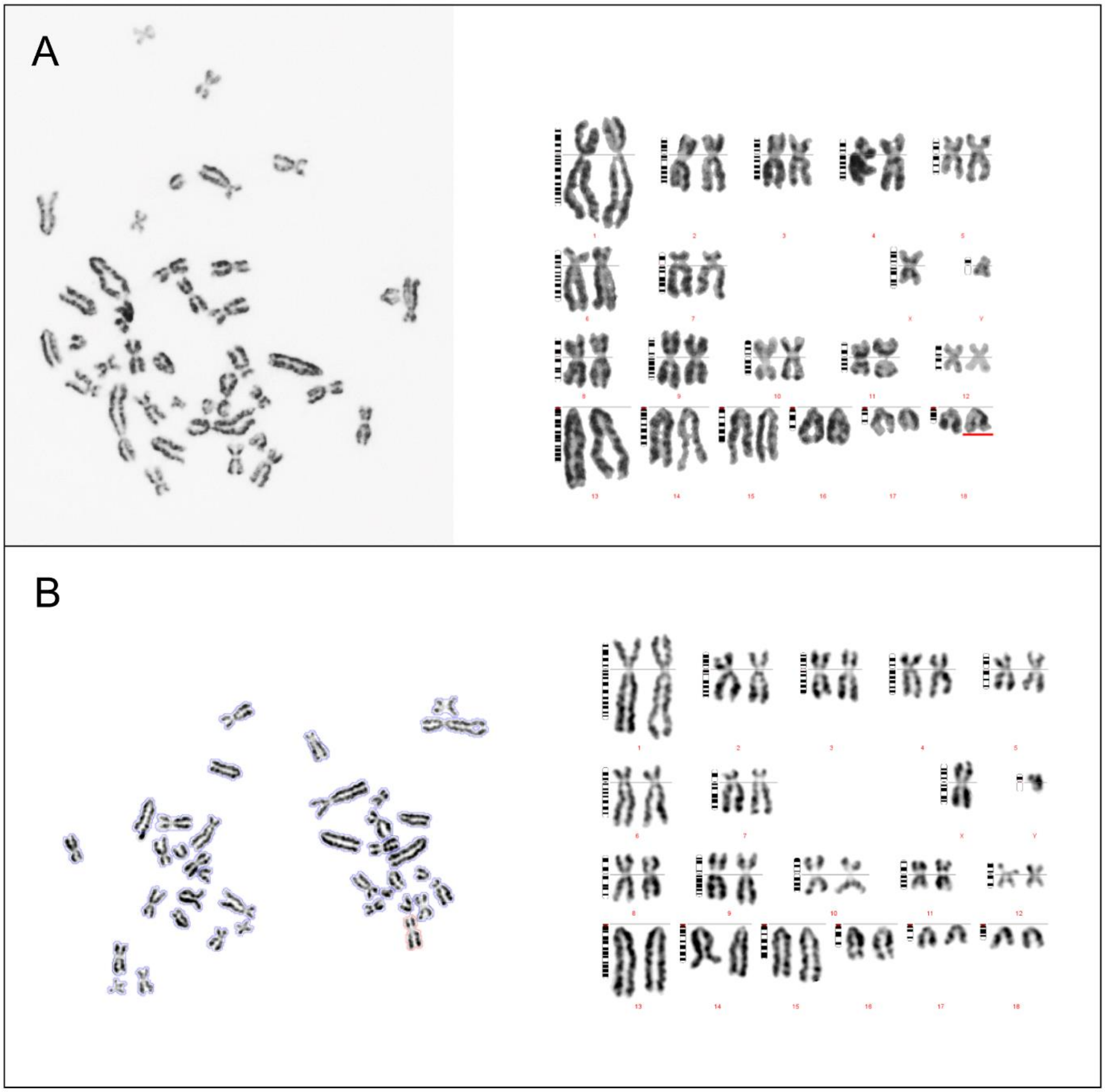


**Fig. S2. Normal karyotypes of 3KO·9TG pigs and PERVKO·3KO·9TG pigs.** 3KO·9TG (A) and PERVKO·3KO·9TG (B) fibroblasts were karyotyped using Giemsa-staining-based G-banding technique. Metaphase spreads were analyzed using SmartType software. Both 3KO·9TG pigs and PERVKO·3KO·9TG pigs show normal [36 + XY] karyotypes.

| Pig | Variant type | gRNA | 1 mismatches | 2 mismatches | 3 mismatches | 4 mismatches |
|---|---|---|---|---|---|---|
| 3KO·9TG | SNV/INDEL | B4GALNT2 | 0 | 0 | 0 | 2[a] |
| | | CMAH | 0 | 0 | 0 | 0 |
| | | GGTA | 0 | 0 | 0 | 0 |
| | SV | B4GALNT2 | 0 | 0 | 0 | 0 |
| | | CMAH | 0 | 0 | 0 | 0 |
| | | GGTA | 0 | 0 | 0 | 0 |
| PERVKO·3KO·9TG | SNV/INDEL | PERV-3N | 0 | 0 | 1[b] | 0 |
| | | PERV-5N | 0 | 1[c] | 0 | 0 |
| | | PERV-9N | 0 | 0 | 0 | 0 |
| | SV | PERV-3N | 0 | 0 | 0 | 0 |
| | | PERV-5N | 0 | 0 | 0 | 0 |
| | | PERV-9N | 0 | 0 | 0 | 0 |

**Fig. S3. Summary of off-target analysis on 3KO·9TG pigs and PERVKO·3KO·9TG pigs.** We detected no structural variant (SV) off-target candidates for 3KO·9TG pigs and PERVKO·3KO·9TG pigs. We detected 2 single nucleotide variant/insertion-deletion (SNV/INDEL) off-target candidates for 3KO·9TG pigs and PERVKO·3KO·9TG pigs respectively ("a": an intergenic insertion and an intronic insertion; "b": an intronic deletion; "c": an intergenic insertion and an intergenic deletion at the same site). None of these confers protein coding change according to VEP annotation v93.3. We called SNV/INDEL using GATK v4.0.7.0 with standard filtering, and SV using 10x Genomics Longranger v2.2.2 with default parameters. We call a variant as an off-target candidate if all the following criteria are met:

(1) It is within 10 bp distance to the PAM of a predicted gRNA off-target site by CRISPRSeek v1.22.1;
(2) The allele frequency difference between the sample and its parental line does not differ significantly from 0.5 using Two-proportion Z-test;
(3) No obvious error by manual check of raw reads on IGV.

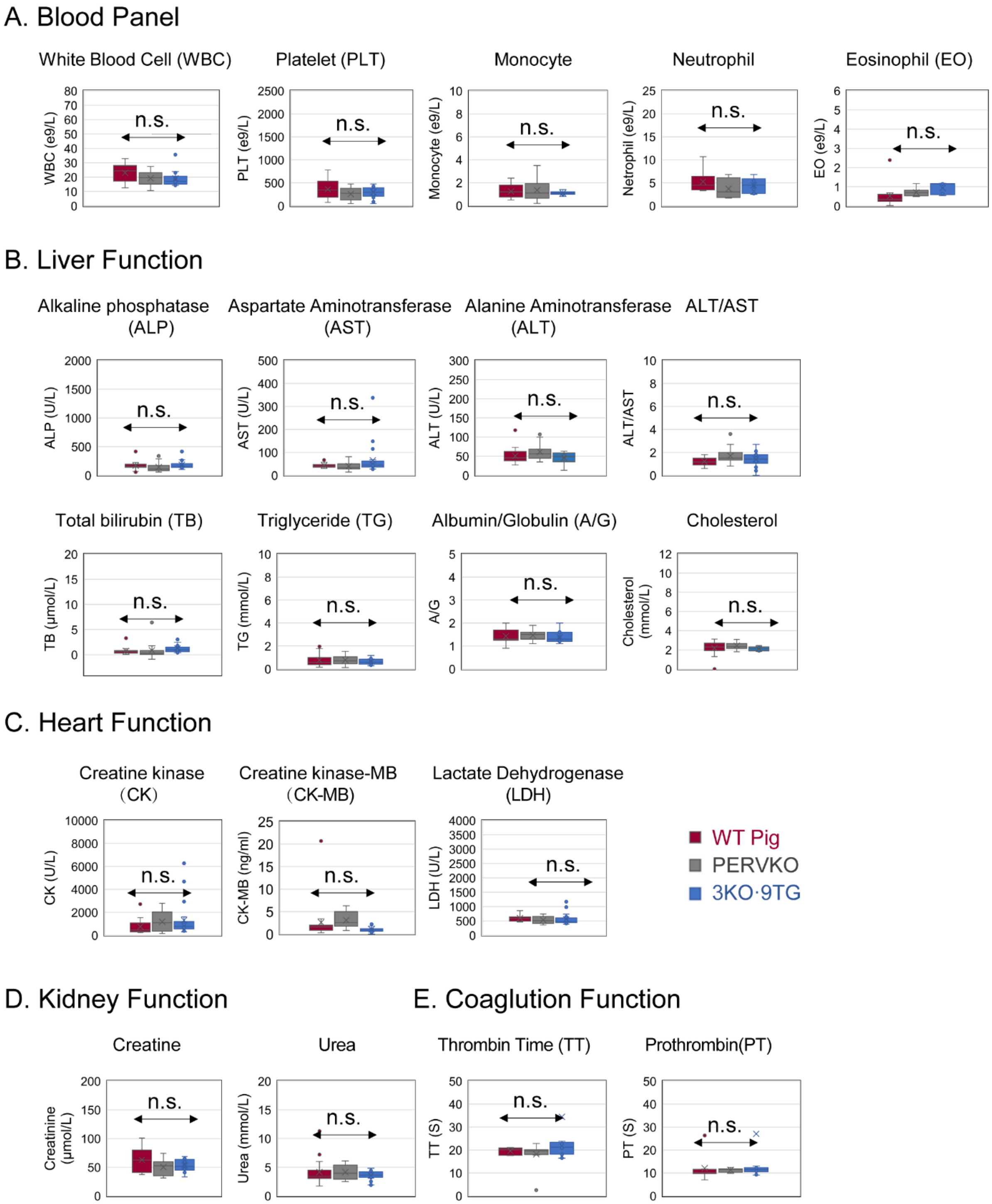


**Fig. S4. PERVKO pigs and 3KO·9TG pigs show similar pathophysiology, compared with WT pigs in parameters measured, including complete blood count (A), liver function (B), heart function (C), kidney function (D), and coagulation function (E).** The sample numbers for PERVKO pigs, 3KO·9TG pigs and WT pigs are 18, 16 and 21, respectively. "no sig" denotes no statistical significance among the PERVKO, 3KO·9TG and WT groups by pairwise T-test analysis.

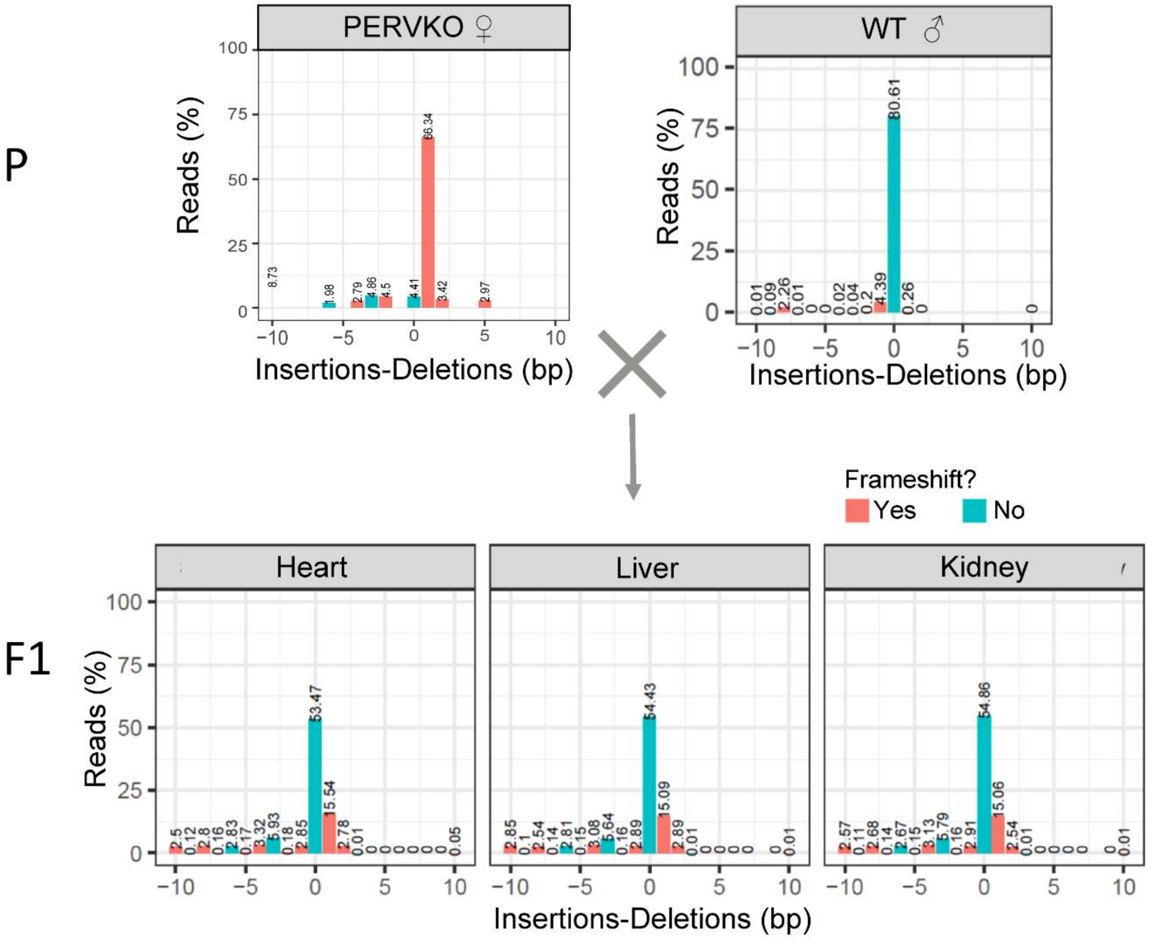


**Fig. S5. The genetic modification of PERVKO can be inherited following Mendelian genetics during natural mating production.** The x-axis represents the total number of shifted bases calculated as the sum of insertions subtracting the sum of deletions. The y-axis represents the percentage of reads. The red and green color indicate frameshift or no frameshift respectively. One PERVKO pig (P) mated with wild type Bama pig and generated 11 piglets (F1). The liver, kidney, and heart tissues of one F1 piglet were analyzed by high-throughput DNA sequencing together with parental fibroblasts to assess the inheritance of the PERVKO modifications. PERVKO has ~96% of PERV alleles to be knockout, while the WT pig has ~80% PERV alleles to be wild type (Insertions-Deletions=0). Of note, some PERV copies in the WT sample may be naturally occurring germline PERVKO alleles. In comparison, the liver, kidney and heart tissues of the F1 piglet has only ~50% PERV copies to carry knockout. The pattern is similar among tissues, indicating that the PERVKO modification is stably inherited following Mendelian genetics.

**A** At the genomic DNA level

| Group | Clone | 9TG | 3KO | | |
|---|---|---|---|---|---|
| | | 9TG | GGTA1 | CMAH | B4GALNT2 |
| | WT Pig | | 0/0 | 0/0 | 0/0/0/0 |
| | 3KO·9TG | √ | -10/9TG-Insert | +2/-391 | -13/-13/-14/-14 |
| | 3KO Pig | | +1/+1 | +1/+1 | -13/-13/-13/-14 |
| 3KO·9TG x WT Offspring | F1-1 | | 0/-10 | 0/-391 | -13/-14/0/0 |
| | F1-2 | √ | 0/9TG-Insert | 0/-391 | -13/-13/0/0 |
| | F1-3 | √ | 0/9TG-Insert | 0/-391 | -13/-13/0/0 |
| | F1-4 | | 0/-10 | 0/-391 | -13/-13/0/0 |
| | F1-5 | √ | 0/9TG-Insert | 0/+2 | -13/-14/0/0 |
| | F1-6 | | 0/-10 | 0/-391 | -13/-13/0/0 |
| | F1-7 | | 0/-10 | 0/-391 | -13/-14/0/0 |
| | F1-8 | | 0/-10 | 0/-391 | -13/-13/0/0 |
| | F1-9 | √ | 0/9TG-Insert | 0/+2 | -13/-13/0/0 |
| 3KO·9TG x 3KO Offspring | F1-1 | √ | +1/9TG-Insert | +1/+2 | -13/-13/-13/-13 |
| | F1-2 | | -10/+1 | +1/-391 | -13/-13/-13/-14 |
| | F1-3 | √ | -10/+1 | +1/+2 | -13/-13/-13/-13 |
| | F1-4 | | -10/+1 | +1/+2 | -13/-13/-13/-13 |
| | F1-5 | √ | +1/9TG-Insert | +1/-391 | -13/-13/-13/-14 |
| | F1-6 | √ | +1/9TG-Insert | +1/+2 | -13/-13/-13/-14 |
| | F1-7 | | -10/+1 | +1/+2 | -13/-13/-13/-14 |
| | F1-8 | | -10/+1 | +1/-391 | -13/-13/-13/-14 |
| | F1-9 | | -10/+1 | +1/-391 | -13/-13/-13/-13 |
| | F1-10 | √ | +1/9TG-Insert | +1/+2 | -13/-13/-13/-14 |
| | F1-11 | | -10/+1 | +1/+2 | -13/-13/-13/-14 |

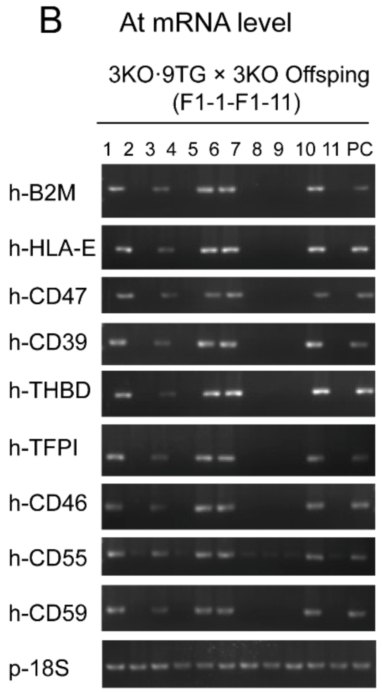


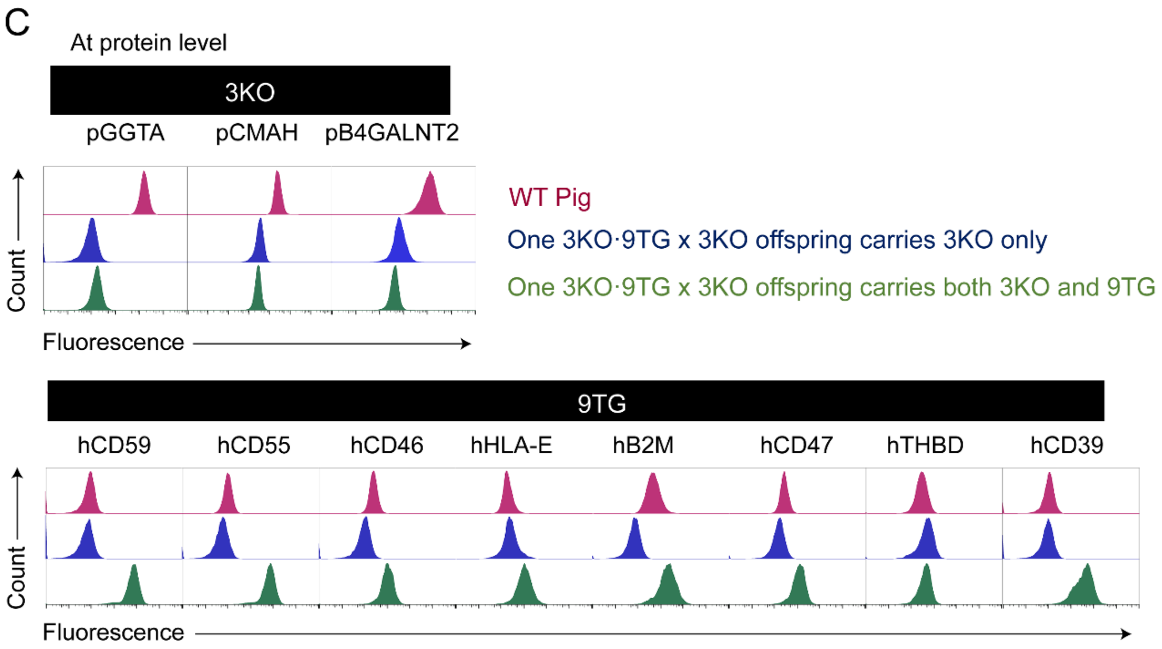


**Fig. S6.** The genetic modifications (3KO and 9TG) of 3KO·9TG pigs can be transmitted to the next generation following Mendelian genetics through natural mating production, as validated at genomic DNA (A), mRNA (B) and protein Level (C). We mated nine WT pigs with 3KO·9TG pigs, and eleven 3KO pigs with 3KO·9TG pigs, and counted the presence of 3KO and 9TG alleles in the F1 progeny. (A) For the 9TG, approximately half of the progeny of 3KO·9TG x WT pigs and 3KO·9TG x 3KO pigs carry the transgenes in the genome. For GGTA1, CMAH and B4GALNT2, the progeny of 3KO·9TG X WT pigs are all heterozygous knockout, and the progeny of 3KO·9TG X 3KO pigs are all homozygous or compound homozygous knockout. Of note,

B4GALNT2 was analyzed as having four alleles because of the inclusion of its highly homologous pseudogene. (B) Approximately half (5/11) of the progeny of 3KO·9TG X 3KO pigs express the mRNA corresponding to the 9TG in their mRNA transcripts. (C) Two example offspring piglets from 3KO·9TG x 3KO, with one carrying 3KO only, and the other carrying both 3KO and 9TG, as verified by FACS.

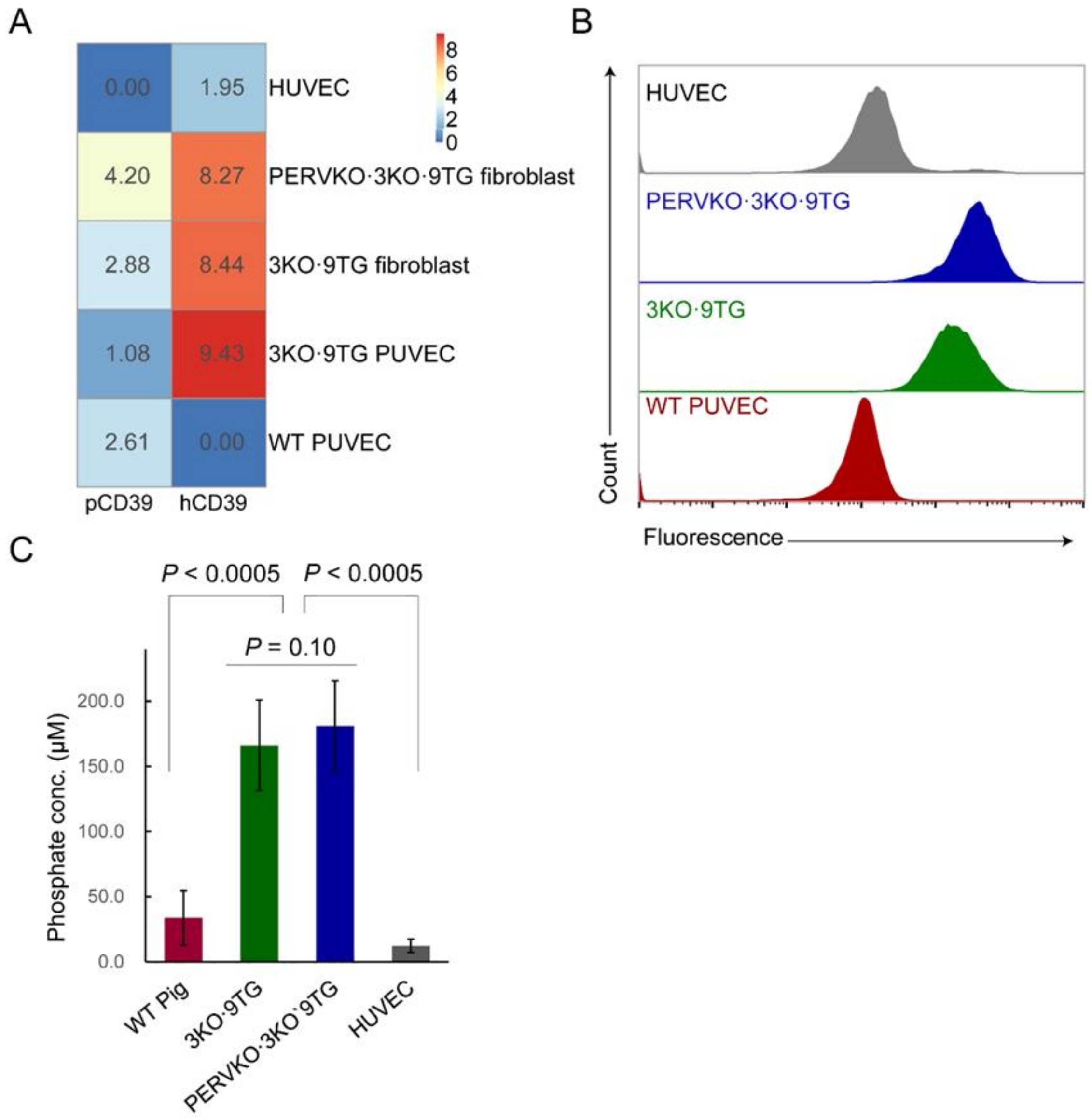


**Fig. S7. 3KO·9TG PUVECs and PERVKO·3KO·9TG PUVECs show significantly higher CD39 ADPase biochemical activity compared to WT PUVECs and HUVECs. (A)** RNA-Seq revealed that the expression level of CD39 in 3KO·9TG PUVECs and PERVKO·3KO·9TG PUVECs is higher than the endogenous level of CD39 in HUVECs (hCD39) and WT PUVECs (pCD39, n = 2). Expression level is color coded in blue-yellow-red to represent low-medium-high and the TPM is shown in each cell. **(B)** FACS revealed that 3KO·9TG PUVECs and PERVKO·3KO·9TG PUVECs have higher human CD39 protein expression than WT PUVECs and HUVECs. **(C)** 3KO·9TG PUVECs and PERVKO·3KO·9TG PUVECs have significantly higher ADPase biochemical activity of CD39 as measured by phosphate production when incubated with ADP. The higher CD39 ADPase biochemical activity is consistent with its higher CD39 protein expression level in 3KO·9TG PUVECs and PERVKO·3KO·9TG PUVECs. Error bars indicate the standard deviation (n = 6).

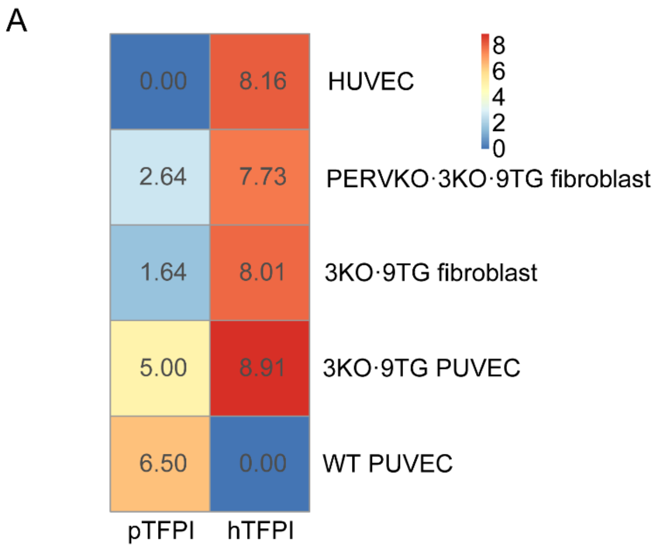


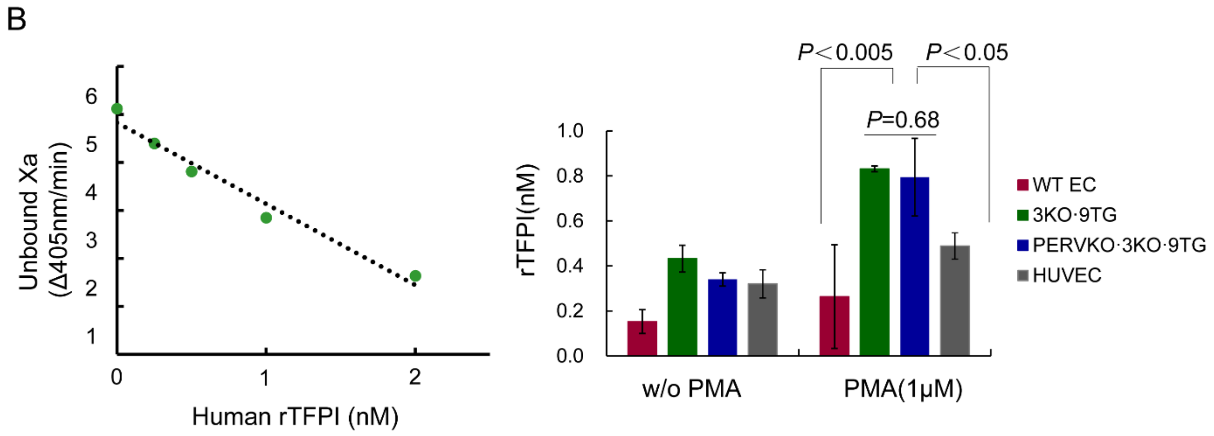


**Fig. S8. Activated 3KO·9TG PUVECs and PERVKO·3KO·9TG PUVECs express human TFPI on cell surface and show significantly higher binding ability to human Xa compared to WT PUVECs and HUVECs *in vitro*. (A)** RNA-Seq revealed that the expression level of TFPI in 3KO·9TG PUVECs is higher than the endogenous level of TFPI in HUVECs (hTFPI) and WT PUVECs (pTFPI, n = 2). Expression level is color coded in blue-yellow-red to represent low-medium-high and the TPM is shown in each cell. **(B)** Activated 3KO·9TG PUVECs and PERVKO·3KO·9TG PUVECs show significantly higher Xa binding ability compared to WT PUVECs and HUVECs *in vitro*. Left panel: Human recombinant TFPI (rTFPI) protein was applied at different concentrations to generate a standard curve between the rTFPI concentration and the unbound Xa level. Right panel: The TFPI level on different PUVECs and HUVECs can be estimated by using the standard curve on the left, which reflects the TFPI binding ability to human Xa. PMA (1 μM): PUVECs and HUVECs were activated by PMA for 6 hours, which leads to the translocation of hTFPI from cytosol to the cell membrane in 3KO·9TG PUVECs and PERVKO·3KO·9TG PUVECs. Error bars indicate the standard deviation (n = 4).
